# Nickel binding to c-Src SH3 domain facilitates crystallization

**DOI:** 10.1101/2025.05.08.652639

**Authors:** Xander Calicdan, Oriana S. Fisher, Byung Hak Ha, Titus J. Boggon, Amy L. Stiegler

**Author notes:** Address correspondence to these authors and.

## Abstract

Numerous X-ray crystal structures of the c-Src SH3 domain have provided a large sampling of atomic-level information for this important signaling domain. Multiple crystal forms have been reported, with variable crystal lattice contacts and chemical crystallization conditions. Here, we report a unique crystal structure of Src SH3 domain in trigonal space group *H*3_2_ to 1.45 Å resolution. Crystal packing and anomalous scattering reveal that this crystal form is mediated by two ordered nickel ions provided by the crystallization buffer. Nickel coordination occurs in a 2:2 stoichiometry which dimerizes two SH3 domain monomers across a pseudo-twofold rotation axis and involves the native N-terminal c-Src SH3 amino acid sequence, a surface-exposed histidine residue, and ordered water molecules. This study provides an example of metal binding by N-terminal protein residues that contrasts the amino terminal copper and nickel binding (ATCUN) motif and is an alternative avenue for crystallization of the Src SH3 domain.

**STRUCTURED ABSTRACT:** *Introduction:* Numerous X-ray crystal structures of the c-Src SH3 domain have provided a large sampling of atomic-level information for this important signaling domain. Multiple crystal forms have been reported, with variable crystal lattice contacts and chemical crystallization conditions.

*Materials and Methods:* We crystallize the c-Src SH3 domain in a crystallization buffer containing NiCl_2_.

*Results:* A unique crystal structure of Src SH3 domain in trigonal space group *H*3_2_ to 1.45 Å resolution is determined. Crystal packing and anomalous scattering reveal that this crystal form is mediated by two ordered nickel ions provided by the crystallization buffer. Nickel coordination occurs in a 2:2 stoichiometry which dimerizes two SH3 domain monomers across a pseudo-twofold rotation axis and involves the native N-terminal c-Src SH3 amino acid sequence, a surface-exposed histidine residue, and ordered water molecules.

*Discussion:* This study provides an example of metal-mediated crystallization and metal binding by N-terminal protein residues that contrasts the amino terminal copper and nickel binding (ATCUN) motif.

**Conclusion:** Alternative avenues for helps widen the potential for future crystallography-based studies of the c-Src SH3 domain.

## 1. INTRODUCTION

Cellular signaling proteins are frequently comprised of multiple modular domains with independent fold and function [1]. Among these protein domains, the Src homology 3 (SH3) domain is one of the most abundant [2]. As its name implies, the SH3 domain was first identified as a conserved region shared among the amino acid sequences of cellular Src (c-Src) tyrosine kinase and the related cellular Abelson tyrosine kinase (c-Abl), phospholipase C, and the adapter protein Crk [3]. In total, 298 SH3 domains were recently reported in 221 human proteins [4]. SH3 domains are also identified in all eukaryotes as well as in some prokaryotes and viruses [4]. These domains typically mediate protein-protein interactions and affect enzyme regulation, substrate binding, protein localization, and motility, regulating pathways that impact many cellular processes including proliferation, migration, endocytosis, and actin cytoskeleton rearrangement (recently reviewed in [5]).

The SH3 domain is a small protein fold comprised of approximately 60 amino acids. The first experimental three dimensional structures of SH3 domains were from the Src Family Kinases (SFKs) Fyn [6] and c-Src [7], and from alpha-spectrin [8]. Together, these structures revealed a conserved SH3 fold comprising two small beta sheets with loops of variable lengths. This fold typically recognizes polyproline-II helical P-x-x-P motifs (or simply ’PxxP’, where x is any amino acid) but can also bind other protein targets in atypical interactions [2, 9–11]. SH3 domain binding to polyproline motifs has been well studied by structural techniques including X-ray crystallography that have collectively helped resolve the binding site, specificity determinants, and bidirectionality of peptide binding [7, 12–15].

The founding SH3 domain was identified in c-Src [3] and has been shown to play a major role in autoinhibition of its kinase activity. In addition to the SH3 domain, SFKs include three other conserved domains: Src homology 1 (SH1) that contains the kinase domain, Src homology 2 (SH2) that binds directly to phosphorylated tyrosine residues, and a short Src homology (SH4) region at the N-terminus which is lipid modified (reviewed in [16]). In the absence of activating signals, intradomain interactions maintain Src in an autoinhibited “closed” conformation; specifically, the SH2 domain binds an inhibitory phosphotyrosine pY-527 in the C-terminal tail, while the SH3 domain binds the linker between SH2 and kinase domains at a non-proline helical-like sequence [17]. In this conformation, the SH3 and SH2 domains impinge on the kinase domain and restrict it to its inactive conformation. In response to signaling, these SH3- and SH2-mediated autoinhibitory restraints are released, and the kinase domain adopts its active conformation. The SH3 domain of SFKs binds protein targets including proline enriched phosphatase (PEP) [18], the adapter protein Crk-II [19], the Nef protein of human immunodeficiency virus-1 (HIV-1) [20], the 3BP1 RhoGAP [21], choline kinase alpha (ChoKα) [22], and beta-arrestin [23].

The three-dimensional structure of the isolated SH3 domain of c-Src has been reported by numerous X-ray crystallographic and NMR studies [24]. Among the X-ray crystal structures available for Src SH3 in the Protein Data Bank, space group identity, unit cell dimensions, and/or lattice contacts are often replicated, presumably owing to the similarity in crystallization conditions or in the sequence of the recombinant protein. Here, we report an X-ray crystal structure of the c-Src SH3 domain in space group *H*3_2_, which has not previously been observed. In the structure we find lattice contacts that are mediated by interchain nickel coordination. This coordination is achieved by the native protein sequence and demonstrates that native amino acids can serve as nickel coordinating residues. This crystal packing arrangement may widen the potential for future crystallography-based studies of the c-Src SH3 domain.

## 2. MATERIALS AND METHODS

### 2.1. Expression construct and expression

Human c-Src (UNIPROT ID: P12931) cDNA encoding the SH3 domain residues 85-143 was PCR-amplified using Platinum *Taq* HiFi polymerase (ThermoFisher) using a forward oligo containing an NdeI restriction site and encoding for a tobacco etch virus (TEV) protease recognition site, and a reverse oligo containing an XhoI restriction site. The PCR product and pET-28a(+) plasmid were separately doubly digested with NdeI/XhoI in CutSmart buffer (New England Biolabs), and ligated together using T4 DNA ligase (New England Biolabs) to yield an expression construct for N-terminal His_6_-tagged protein with both thrombin and TEV protease recognition sequences. A positive clone was verified by whole-plasmid sequencing (Plasmidsaurus). See Table S1. Protein expression was carried out in Rosetta(DE3) *E. coli* cells transformed with the expression plasmid. Cells were cultured in the presence of 25 µg/ml kanamycin and 17 µg/ml chloramphenicol at 37°C shaking at 200 rpm in Miller’s LB Broth (Fisher Scientific) to an OD_600_ of 0.8, then cooled to 18°C prior to induction of protein expression with 0.2 mM Isopropyl β-D-1-thiogalactopyranoside (IPTG). Expression was conducted overnight (approximately 20 hours) at 18°C shaking at 200 rpm. Cells were harvested by centrifugation.

### 2.2. Protein purification

Cells were resuspended in lysis buffer (500 mM NaCl and 50 mM HEPES pH 7.5) and flash frozen in a dry ice/ethanol bath. The sample was thawed and lysozyme added to a final concentration of 0.1 mg/ml, followed by two more cycles of freeze/thaw. The sample was lysed by sonication for 3 cycles of 2 minutes using 10 second on/off pulses at 35 % with a half-inch probe (Q500, QSonica). DNase I (bovine pancrease, GoldBio) was added to 10 Units/ml and incubated 10 min on ice. The total lysate was clarified by centrifugation for 1 hr at 4°C at 32,928 x *g*. Clarified lysate was applied to 1 ml Ni-NTA agarose beads (Qiagen) in a sealed gravity flow column and incubated rocking for 1 hour at 4°C. The flow-through sample was collected, and beads washed with wash buffer (lysis buffer containing 20 mM imidazole). Recombinant Tobacco Etch Virus (rTEV) protease was added to the bead-bound His_6_-tagged Src SH3 protein and incubated overnight at 4°C. The liberated untagged c-Src SH3 domain protein was then collected off the bead by washes with 1 ml aliquot of wash buffer and further purified by size exclusion chromatography (Superdex 75 Increase 10/300 GL) in buffer containing 150 mM NaCl, 20 mM Tris pH 7.5. The purified protein was concentrated in a spin concentrator (Amicon Ultra 3 kDa MWCO, Millipore Sigma) to a final concentration of 11 mg/ml. Protein concentration was determined by NanoDrop (Thermo Fisher) A280 nm blanked with size exclusion buffer using an extinction coefficient of 16,960 M^-1^ cm^-1^ calculated using the primary amino acid sequence of the protein.

### 2.3. Protein crystallization

Purified c-Src SH3 domain protein at 11 mg/ml was crystallized in hanging drops by vapor diffusion at room temperature in 24-well VDX plates (Hampton Research) over 500 µl of reservoir solution. Crystallization drops contained 1 µl of protein with 1 µl of reservoir buffer containing 5 mM Nickel chloride, 1.7 M Ammonium Sulfate, 10 % glycerol, and 0.1 M HEPES pH 7.5. Crystals grew with morphology of rectangular prisms with typical edge dimension between 25 to 100 µm and appeared after 1 to 3 days. Crystals were harvested, washed in cryopreservative solution containing reservoir solution and 20% glycerol, and flash cooled in liquid nitrogen by manual plunging. See Table S2.

### 2.4. X-ray data collection and processing

X-ray diffraction data were collected at Brookhaven National Laboratory National Synchrotron Light Source II beamline 17-ID-2 (FMX) [25]. Crystal mounting, raster scanning, centering, and data collection were each performed automatically. Data were collected at wavelength 0.979315 Å. Autoprocessing was performed in the autoPROC pipeline [26] which utilizes XDS for indexing and integration [27]. Individual datasets from a single crystal contain 1200 images (4800 images in total) with 0.01 sec exposure and oscillation range of 0.2° per image for a total rotation range of 240°. Reflection data from 4 individual crystals were input to Aimless for scaling and merging [28]. An anomalous signal was automatically detected by XDS in each of the 4 datasets; thus, Friedel pairs were treated separately. Both intensities (*I*(+) and *I*(-)) and structure factors (*F*(+) and *F*(-)) were output to MTZ. A resolution cutoff of 1.45 Å was determined based on a combination of values of *I*/σ*I* > 1.5, CC½ > 0.5 and *R*_pim_ < 0.5, using the guidelines of [28]. Crystals belong to space group *H*3_2_ with unit cell dimensions a = 63.77 Å, b = 63.77 Å, c = 271.71 Å, α = β = 90°, γ = 120°. Phenix Xtriage analysis [29] reveals that the diffraction data contain an off-origin native Patterson peak of height 31% due to translational non-crystallographic symmetry (tNCS). See Table 1 and Table S3.

**Table 1.** X-ray diffraction data collection and processing statistics.

|  |  |
| --- | --- |
| Wavelength (Å) | 0.979315 |
| Number of crystals merged | 4 |
| Resolution range (Å) | 31.95 - 1.45 (1.48 - 1.45) |
| Anomalous resolution limit (Å)<sup>+</sup> | 2.00 |
| Space group | *H*3<sub>2</sub> |
| Cell dimensions<br>a, b, c (Å)<br>$\alpha$, $\beta$, $\gamma$ (°) | 63.77, 63.77, 271.71<br>90, 90, 120 |
| Number of reflections |  |
| Total observations | 2,030,203 |
| Unique (anomalous) | 73,036 |
| Unique (non-anomalous) | 38,628 |
| Completeness (%) | 100 (100) |
| Anomalous completeness (%) | 100 (100) |
| Redundancy | 52.6 (46.8) |
| Anomalous redundancy | 27.2 (23.8) |
| Mean $I/\sigma(I)$ | 22.9 (1.8) |
| Anomalous Mean $dI/\sigma(dI)$ | 1.3 (0.89) |
| Mosaicity (°) of merged data | 0.07 |
| $CC_{1/2}$ | 1.00 (0.784) |
| $R_{pim}$ | 0.012 (0.509) |
| Wilson $B$-factor (Å<sup>2</sup>) | 23.07 |

### 2.5. Structure determination, model building and refinement

The crystal structure was determined by molecular replacement in Phaser [30] using as the search model a previously published crystal structure of chicken c-Src SH3 domain truncated to residues 86-140 and lacking hydrogen atoms (PDB: 4JZ4 chain A [31]). Translational NCS (tNCS) was detected automatically and applied during Phaser with NMOL equal to 2 and an inter molecule vector of (1, 0, 0.5). A molecular replacement solution containing 4 copies of the SH3 domain was found with top TFZ score of 10.9 and eLLG of 1070, initial *R*_work_ = 30.2% and *R*_free_ = 32.1%. Autobuilding was performed by Phenix autobuild [32] to 227 residues in 4 chains and 169 waters with *R*_work_ = 25.2% and *R*_free_ = 26.53%. Manual model building was performed in Coot [33] and refinement in Phenix [29] using occupancy refinement and anisotropic *B*-factor refinement of protein and Nickel atoms. Waters were built automatically during autobuild and refinement, refined with isotropic *B*-factors, and were inspected and edited manually in Coot. Glycerol molecules from the cryopreservative solution were modeled manually and refined with isotropic *B*-factors. Nickel atoms were modeled into anomalous difference density. Secondary structure definitions are obtained by DSSP [34]. The free set for *R*_free_ calculation was randomly selected and flagged as 5% of reflections using Phenix reflection file editor. See Table 2.

**Table 2.** Structure refinement.

|  |  |
| --- | --- |
| Resolution range (Å) | 31.89 - 1.45 (1.47 - 1.45) |
| Completeness (%) | 99.9 (100) |
| No. of reflections, working set | 73,036 (2,734) |
| No. of reflections, test set | 3,450 (137) |
| Reflections, test set size (%) | 4.72 (5.0) |
| Final $R_{\text{work}}$ | 16.25 (27.88) |
| Final $R_{\text{free}}$ | 19.61 (37.45) |
| No. of non-H atoms |  |
| Protein | 1,888 |
| Metals (Ni<sup>2+</sup>) | 2 |
| Waters, other solvent | 163, 54 |
| Total | 2,107 |
| R.M.S. deviations from ideality* |  |
| Bonds (Å) | 0.005 |
| Angles (°) | 0.879 |
| Average $B$-factors (Å<sup>2</sup>) | |
| Protein | 30.70 |
| Metals (Ni<sup>2+</sup>) | 24.62 |
| Waters, other solvent | 38.87, 58.29 |
| Overall | 32.07 |
| MolProbity Statistics |  |
| Ramachandran plot |  |
| Favored regions (%) | 98.19 |
| Allowed regions (%) | 1.81 |
| Outliers (%) | 0.00 |
| Clashscore | 2.63 |
| Unmodelled residues (%) | 2.12% (5 out of 236 total residues) |
| PDB Code | 9OFX |

Values given in parentheses are for the highest resolution shell.
\* Phenix Geometry restraints library

Phased anomalous difference maps were calculated in the Phenix utility phenix.find_peaks_holes [29] using the protein model phases combined with the structure factors *F*(+) and *F*(-) from the diffraction data. Two strong anomalous peaks of heights 58.9 σ and 43.5 σ were used to guide modeling of two nickel atoms. These Ni^2+^ sites were validated in the CheckMyMetal server [35] and statistics given in Table 3. Refinement of the anomalous scattering factor *f*’’ was calculated in Phenix to a value of 1.745 e-(analysis as per [36] and [37]), which is consistent with the theoretical *f*’’ value of 1.943 e-for Ni at wavelength 0.9793 Å (12,660 eV) obtained via X-ray absorption edge tabulations at http://www.bmsc.washington.edu/scatter/AS_index.html and [38]. A third anomalous peak of 6.9 σ was present; however, a Ni atom was modeled and refined to occupancy of less than 0.2 with high *B*-factors and poor coordination. Therefore, this third nickel ion was omitted from the final model. Structures are illustrated with CCP4mg [39] and ChimeraX [40]. All crystallography software was compiled by SBGrid [41].

**Table 3.** Nickel site information.

|  |  |  |
| --- | --- | --- |
| Anomalous scatterer | Nickel |  |
| Photon energy (eV) | 12,660 |  |
| Theoretical $f''$ (e-) | 1.943 | |
| Refined $f''$ (e-) | 1.745 | |
| Nickel sites | Ni-1 | Ni-2 |
| Peak height ($\sigma$) * | 58.9 | 43.5 |
| CheckMyMetal<sup>+</sup> statistics |  |  |
| $B$-factor ($\text{\AA}^2$) | 22.0 | 27.2 |
| Valence | 2.7 | 2.8 |
| Geometry | Octahedral | Octahedral |
| R.M.S.D. Geometry<sup>+,#</sup> |  |  |
| Angles ($^\circ$) | 4.3 | 6.0 |
| Bonds ($\text{\AA}$) | 0.178 | 0.250 |
| Metal-ligand distances ($\text{\AA}$) | | |
| Gly-85 N | 1.97 | 1.97 |
| Gly-85 O | 1.97 | 1.98 |
| His-125 N (Nε2) | 1.99 | 1.99 |
| Water | 1.97 (1) | 1.97 (4) |
| Water | 1.98 (2) | 1.97 (5) |
| Water | 1.99 (3) | 1.97 (6) |

\* Anomalous peak heights ( $\sigma$ ) calculated in Phenix (phenix.find\_peaks\_holes)
<sup>+</sup> CheckMyMetal Server [35]
<sup>#</sup> FindGeo Server [45]

## 3. RESULTS

### 3.1. Crystallization and structure determination

We set out to reproduce published crystallization conditions curated in the Protein Data Bank for the isolated c-Src SH3 domain protein for didactic purposes. Among those tested, crystallization conditions containing a cocktail of 1.4 - 1.8 M ammonium sulfate, HEPES pH 7.5, and 5 mM NiCl_2_ in the reservoir solution yielded single crystals with rectangular prism morphology with edge length averaging between 25 - 100 µm despite heavy precipitate also being observed in the drops (Figure 1A, Table S2). We observed that when NiCl_2_ was omitted from crystallization conditions, crystal growth was not achieved. Crystals were harvested from the drop, washed in reservoir solution, cryopreserved, and cryocooled in loops for X-ray data collection at the FMX beamline at NSLS-II. Datasets from four individual crystals were merged to achieve a multiplicity of 52.6. Consequently, anomalous signal improved and was detected automatically during data processing (Table 1 and Table S3).

**Figure 1.**
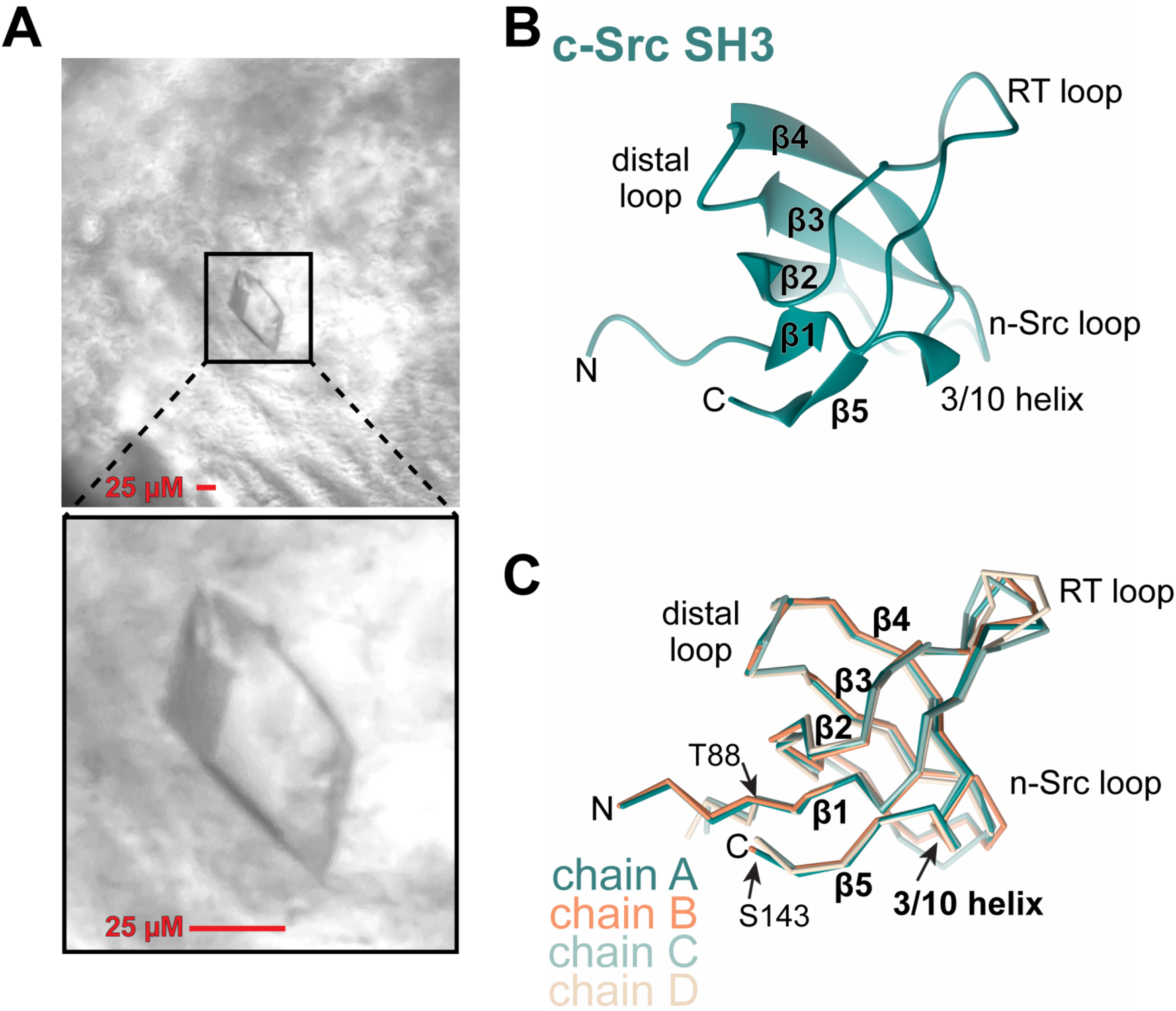
**Crystal structure of c-Src SH3 domain in *H*3_2_ crystal form.** (A) Representative crystals of c-Src SH3 domain grown for this study. A scale bar representing 25 µM is included in each panel. The average crystal has edge dimension of 25 - 100 µm and fit within a 0.05-0.1 mm CryoLoop (Hampton Research). (B) Ribbon diagram of the c-Src SH3 domain crystal structure from this study. Shown is chain A. (C) Superposition of the structures of each chain of c-Src SH3 domain from this study. Shown is the Cα trace of each chain colored dark teal (chain A), orange (chain B), light teal (chain C) and tan (chain D). The positions of Thr-88 and Ser-143 are indicated with arrows. In (B) and (C), the secondary structure elements, loops, and amino- and carboxy-termini are named and labelled.

The structure of the c-Src SH3 domain was determined to 1.45 Å resolution by molecular replacement (Table 2). Crystals belong to space group *H*3_2_ with unit cell dimensions a = b = 63.77 Å, c = 271.71 Å, α = β = 90°, γ = 120° and four molecules per asymmetric unit. The *H*3_2_ space group differs from the anticipated space groups *P*2_1_2_1_2_1_ (PDB: 4jz4 and 4omo [31] and 4rtz, unpublished)) or *P*2_1_ (PDB: 4rtx, unpublished) of the previously reported c-Src SH3 crystals grown in similar NiCl_2_-containing crystallization conditions. Examination of the Protein Data Bank reveals that no available crystal structure of the Src SH3 domain adopts the *H*3_2_ space group or unit cell dimensions of our crystal form. Furthermore, interface search performed in the ’Protein interfaces, surfaces and assemblies’ (PISA) server [42] of crystal lattice contacts in the *H*3_2_ crystal form reveals no common lattice contact in any available crystal structure of c-Src SH3 domain including the *P*2_1_2_1_2_1_ or *P*2_1_ crystal forms. Thus, this previously reported crystallization condition yielded a unique crystal lattice for c-Src SH3.

### 3.2. SH3 domain structure

We performed model building and refinement of the c-Src SH3 domain structure in space group *H*3_2_. As our structure confirms, the c-Src SH3 domain structure is comprised of two antiparallel β-sheets formed by strands β1 - β2 - β5 and strands β3 - β4. These strands are connected by the long RT-loop (β1 - β2), n-Src loop (β2 - β3), distal loop (β3 - β4), and a 3/10 helix (β4 - β5) (Figure 1B and 1C, using nomenclature according to [6]). Chains A, B, and C are fully modeled from residues 85-143, while chain D lacks strong electron density for residues 115-119 (in the n-Src loop) which remain unmodelled. The four copies align closely with one another: upon superposition of residues Thr-88 through Ser-143, the structural alignment results in R.M.S.D. values of 0.38 - 0.99 Å over 54 Cα positions (Figure 1C, [43]). Small positional changes in the RT and n-Src loops are the most pronounced differences between the modelled chains. Comparison of the four copies in the asymmetric unit also revealed differences in the N-terminal residues Gly-85, Val- 86 and Thr-87. These variations in conformation are due to nickel binding which is discussed in detail below, and explain why the asymmetric unit is composed of four individual non-symmetric chains. Our c-Src SH3 structure also closely aligns with previously determined structures, with R.M.S.D. values ranging from 0.4 to 1.5 Å over 54 - 59 Cα positions for many of the available Src SH3 domain structures in the PDB (Figure S1, Dali server [44]). We therefore consider the unique lattice contacts in our *H*3_2_ crystal form are not attributed to large conformational changes or structural rearrangements in the Src SH3 domain protein.

### 3.3. Nickel coordination in the asymmetric unit

Since a strong anomalous signal was detected in the X-ray diffraction data (Table 1 and Table S3), we computed phased anomalous difference maps which revealed two large anomalous peaks with heights of 58.9 σ and 43.5 σ (Figure 2A and Table 3) at locations where auto building had initially modeled water molecules; however, after refinement of these waters, the positive *F*_o_-*F*_c_ difference map remained highly elevated (not shown), suggestive of the presence of a non-water atom (such as a metal) at these positions. As 5 mM NiCl_2_ was required for crystal growth (Table S2) and nickel has a sufficiently high theoretical anomalous *f*’’ atomic absorption value of 1.951 [38] electrons at the X-ray collection wavelength (0.98 Å), we modeled and refined nickel atoms at these anomalous peak positions (Figure 2B). The two nickel sites are located at the interface of Src SH3 chains A and B in a 2:2 stoichiometry. The nickel atoms bridge the A and B chains by coordination of the nitrogen and carboxyl oxygen atoms of the N-terminal residue Gly-85 of one chain and the imidazole epsilon nitrogen atom of His-125 on β3 in the neighboring chain. The crystallization pH value of 7.5 is favorable for metal coordination by histidine since its sidechain, with pKa value of 6.0, will be at least partially deprotonated. Octahedral metal coordination is completed by three water molecules (Figure 2B). The refined nickel-ligand bond distances (1.97 - 1.99 Å) are within the expected range (Table 3 and Figure S2A and S2B). Both nickel sites are validated by CheckMyMetal Binding Site Validation [35] and FindGeo [45] servers (Table 3).

**Figure 2.**
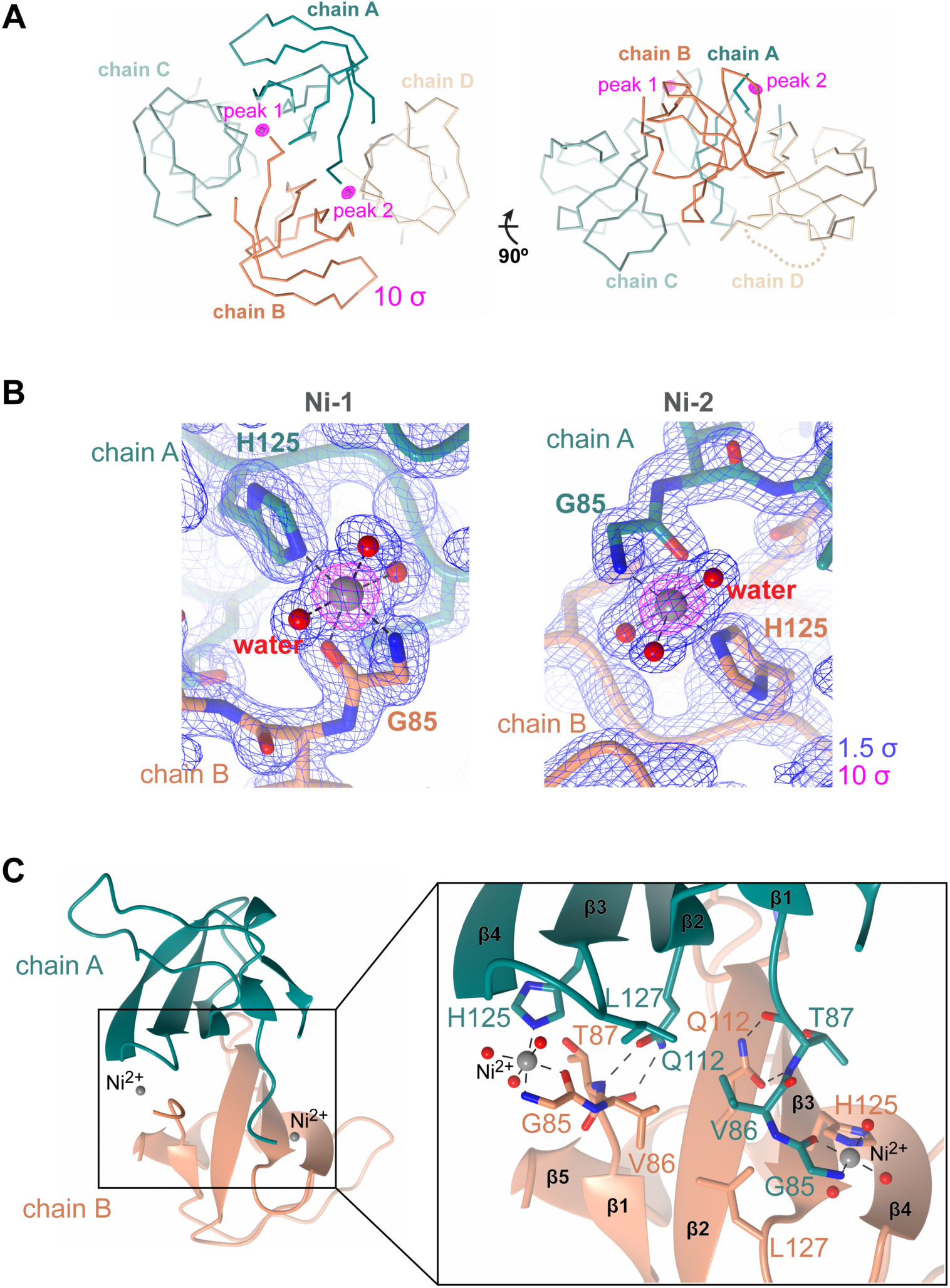
**Nickel coordination by the c-Src SH3 domain.** (A) The asymmetric unit containing four chains of c-Src SH3 domain (Cα traces) and phased anomalous difference map contoured to 10 σ (pink) shows two strong peaks (peak 1 and peak 2) arranged between chains A (dark teal) and B (orange). The two views are related by a 90° rotation about the x-axis away from the reader. (B) Final refined 2*F*_o_-*F*_c_ map (blue, contoured to 1.5 σ) and phased anomalous difference map (pink, contoured to 10 σ). Ni^2+^ atoms are drawn as grey spheres, and water molecules are red spheres. (C) Details of nickel binding. Left panel: overview of chain A/chain B interface. Zoom-in: details of the interface including intrachain nickel coordination (black dashed lines) by Gly-85 and His-125.

We further examined the 2:2 stoichiometry interaction between the Ni^2+^ ions and chains A and B of the SH3 domain. These chains are related by non-crystallographic twofold rotational symmetry and the interface buries a surface area of approximately 440 Å^2^ per chain. In addition to the intrachain Ni^2+^ coordination, the interface includes hydrogen bonding between Gln-112 and Thr-87 and van der Waals interactions including Val-86 and Leu-127 near the nickel binding site (Figure 2C). There is also a small patch of van der Waals interactions involving Asn-115 and Thr-117 on the opposite side of the chains A/B interface (Figure S3). The nickel-mediated symmetrization aids in lattice formation by stabilizing the interactions between chains A and B in the asymmetric unit.

### 3.4. Role of nickel in lattice formation in *H*3_2_ space group

We then more closely examined the other crystal contacts in this *H*3_2_ c-Src SH3 crystal form. The four molecules in the asymmetric unit form two types of equivalent dimers: one between chains A/C and B/D which buries approximately 500 Å^2^ per monomer, and a second between chains A/D and B/C which buries an average of 380 Å^2^ per monomer (Figure 3A). Each of these dimer sets are related by non-crystallographic two-fold rotational symmetry. An interface search of the available structures in the Protein Data Bank for each of these dimers using the PISA server [42] reveals no equivalent Src SH3 interface; thus, the lattice formation in the *H*3_2_ space group is unique among available crystal structures of the Src SH3 domain. The binding of Ni^2+^ across the interface of chains A and B bridges these dimer pairs to help stabilize the *H*3_2_ crystal lattice. Next, examination of the global crystal packing within the unit cell reveals that Ni coordination aides in asymmetric unit formation but is not involved in packing of the asymmetric units into the unit cell (Figure 3B). Taken together, these observations support the conclusion that nickel aids in formation of the asymmetric unit.

**Figure 3.**
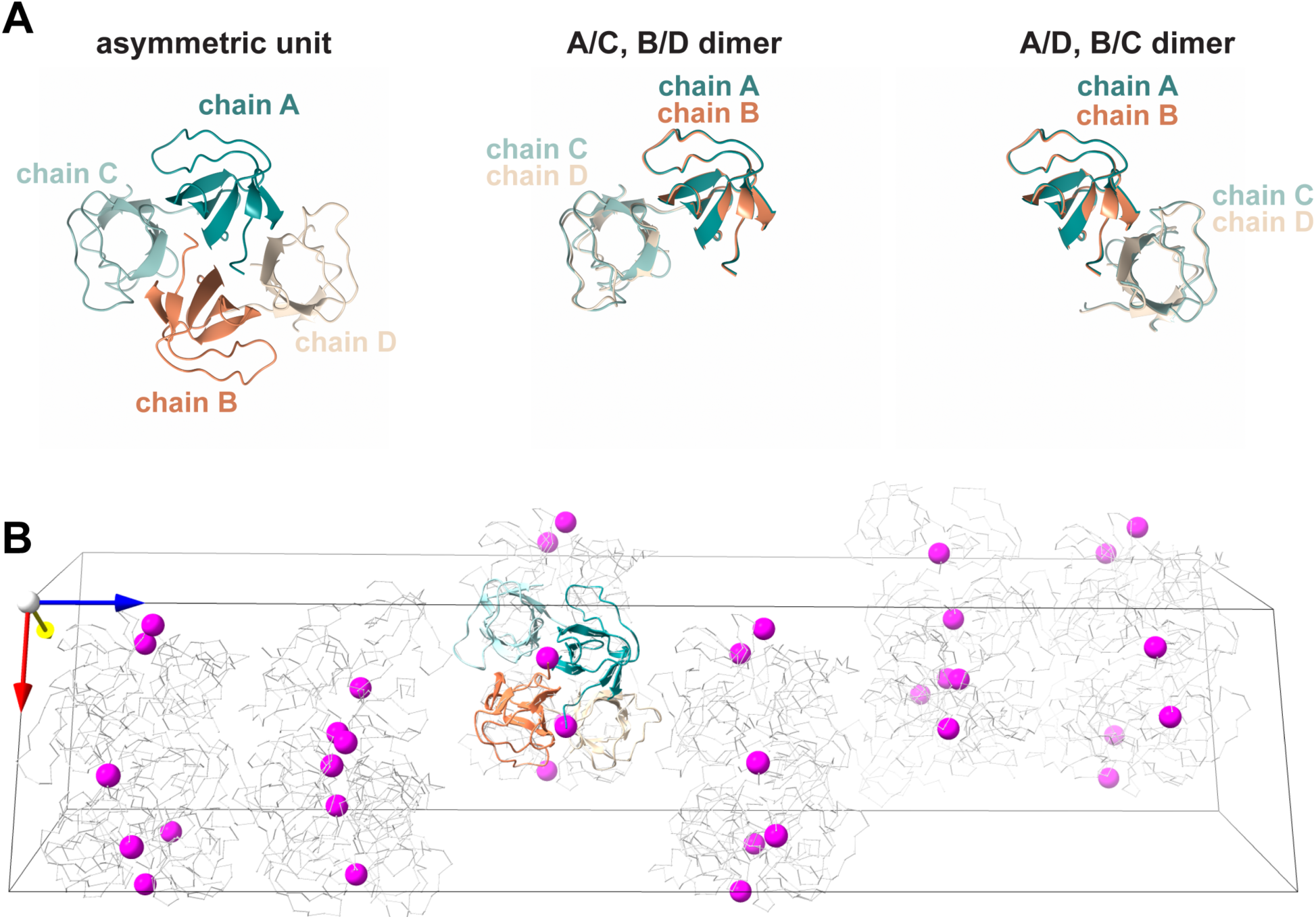
**Lattice contacts in the c-Src SH3 *H*3_2_ crystal form.** (A) Left panel: asymmetric unit arrangement of four copies of the c-Src SH3 domain. Two types of dimers are depicted in the center and right subpanels. (B) Expanded view of the unit cell, which is drawn as a box. The nickel sites are indicated as pink spheres. One copy of the asymmetric unit is drawn with protein ribbons and colored as in part (A). The other protein monomers are drawn as gray Cα traces. The orientation axes x (red), y (yellow) and z (blue) are indicated. The nickel sites bridge monomers A and B in the asymmetric unit but do not further aid in crystal packing between asymmetric units.

To further study this phenomenon, we attempted to detect nickel-induced dimer formation of the SH3 domain by comparing the elution profile from size exclusion chromatography and migration through semi-native SDS-PAGE [46, 47] in the absence and presence of a five-fold molar excess of Ni^2+^ from NiCl_2_; however, neither method yielded conclusive results. Therefore, whether nickel-induced dimerization occurs *in vitro* prior to crystallization or *in situ* in the crystallization dop during vapor diffusion remains unresolved.

### 3.5. Comparison with the amino-terminal copper and nickel (ATCUN) binding motif

The *H*3_2_ crystal form of c-Src SH3 domain differs from those previously reported despite growing in similar crystallization conditions containing NiCl_2_. To understand this discrepancy, we examined the protein constructs used for crystallization. We designed the expression plasmid to encode for a TEV protease recognition sequence (Glu-Asn-Leu-Tyr-Phe-Gln-Gly) occurring C-terminal to the His_6_-tag which is absent from the parental pET-28 vector. After TEV proteolysis of the purified protein, which occurs between the Gln-Gly peptide bond of the recognition sequence, the resulting N-terminus bears the sequence Gly-Val, which is equivalent to Gly-85 and Val-86 in the native Src sequence (Table S1). This cloning scheme yields an all-native amino acid sequence following TEV proteolysis and facilitates the Ni^2+^ ion-mediated crystal form (Figure 2B and 2C).

We next compared the Ni^2+^ binding by c-Src SH3 in the *H*3_2_ space group with the Ni^2+^ binding in the previously reported crystal structures of Src SH3 from similar NiCl_2_-containing crystallization conditions and at similar pH values of 7.5 (PDB: 4jz4 and 4omo [31], 4rtz (unpublished) and 4rtx (unpublished)). In these structures, Ni^2+^ ions are modelled and refined, and each chain of c-Src SH3 coordinates a single Ni^2+^ ion via the N-terminal residues Gly-Ser-His, which form a so-called ’amino terminal copper and nickel’ (ATCUN) binding motif [48]. The classic monomeric ATCUN motif is comprised of the amino-terminal tripeptide consensus sequence (H_2_N)-x-x-His (where x is any amino acid and histidine is at position 3), which is found in the naturally occuring ATCUN site in proteins including human and bovine serum albumin where it is used for copper transport in the blood [49]. The ATCUN motif is also used as an engineered short peptide motif well-suited for metal-coordination (reviewed in [50]) that can chelate divalent metals at very high affinities (100 fM [51]). In the bona fide ATCUN-motif containing Src SH3 protein structures, the nickel ion is coordinated by the ATCUN motif of a single chain in distorted square planar geometry via four-nitrogen chelation from the backbone nitrogen atoms of Gly-81, Ser-82 and His-83 and an imidazole nitrogen from the sidechain of His-83 with bond distances ranging from 1.83 - 2.02 Å (Figure 4A and Figure S2C and S2D). This Gly-Ser-His motif is present at the N-terminus of these c-Src SH3 domain proteins as the result of cloning artifacts and thrombin protease cleavage, and for c-Src SH3 is a non-native amino acid sequence. This differs from our interchain nickel coordination mediated by native amino acid sequences at the N-terminus Gly-85 and Val-86 and the sidechain of His-125 across the dimer interface (Figures 2 and S2A and S2B). Thus, nickel coordination by Src SH3 contrasts binding of metals by the ATCUN motif-containing proteins.

**Figure 4.**
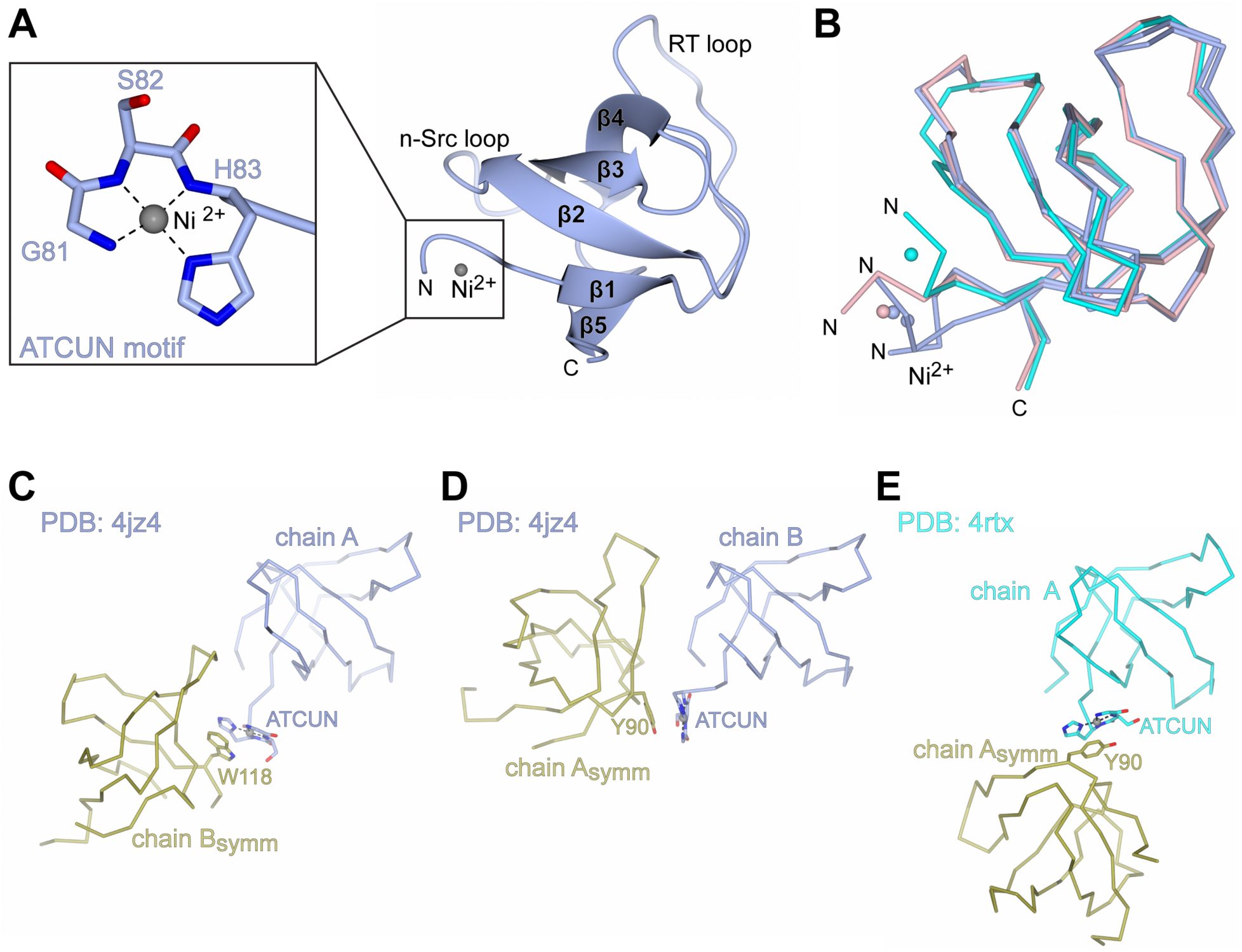
**Nickel coordination by c-Src SH3 domains containing an ACTUN motif.** (A) The ATCUN motif Gly-Ser-His at the N-terminus of Src SH3 domain protein (PDB: 4jz4, [31]) coordinates nickel in square planar geometry. (B) Superposition of c-Src SH3 domains containing ATCUN and coordinating nickel. Included are PDBs 4jz4 (light blue, chains A and B, [31]), 4rtz (light pink, *P*2_1_2_1_2_1_, unpublished) and 4rtx (cyan, *P*2_1_, unpublished). PDB 4omo [31] is equivalent to 4jz4 and is not shown. The nickel sites are drawn as spheres and colored according to the corresponding PDB. (C) Symmetry-related π-stacking interaction involving the nickel-binding ATCUN motif of chain A (light blue) and the sidechain of Tyr-90 in symmetry-related chain B (chain B_symm_, gold). PDB: 4jz4 [31]. (D) Symmetry-related π-stacking interaction involving the nickel-binding ATCUN motif of chain B (light blue) and the sidechain of Trp-118 in a symmetry-related chain A (chain A_symm_, gold). PDB: 4jz4 [31]. (E) Symmetry-related π-stacking interaction involving the nickel-binding ATCUN motif of one chain (cyan) and the sidechain of Tyr-90 in a second chain (gold). PDB: 4rtx (unpublished). For comparison, the orientation of the main SH3 domain molecule is equivalent in parts C, D, and E.

Despite coordinating nickel in a distinct way than we observe here, the ATCUN motifs in the previous nickel-containing Src SH3 domain crystal structures still aided in crystal packing. Among these ATCUN structures are different N-terminal conformations with flexibility in the positioning of the Ni-bound ATCUN motifs with respect to the folded portion of the SH3 domain (Figure 4B). In addition, these Src SH3 crystals adopt different space groups (*P*2_1_2_1_2_1_ and *P*2_1_) and in the case of *P*2_1_2_1_2_1_, two different unit cells with different packing interfaces. It has been postulated that this N-terminal flexibility contributes to different crystal packing among these earlier structures [52]. In contrast to the *H*3_2_ crystal form, the previous Ni-bound ATCUN motifs do not bridge monomers by direct nickel coordination, rather, the ATCUN motif promotes crystal packing by forming π-stacking interactions with one of two aromatic residues in neighboring symmetry-related chains, either Trp-118 (Figure 4C) or Phe-90 (Figure 4D and 4E). The fourth

ATCUN-containing SH3 structure, PDB 4rtz (unpublished), is a co-crystal structure with a PxxP peptide. In this crystal form, the peptide contributes to crystal packing and the ATCUN-Ni participates in minor van der Waals interactions with a neighboring chain yet does not undergo π-stacking (not shown). Thus, flexibility in the N-terminal ATCUN motifs promote different crystal packing. The *H*3_2_ crystal form represents further exploration of Src SH3 N-terminal mediated nickel coordination.

### 3.6. The Src SH3 *H*3_2_ crystal form is not compatible with binding extended PxxP peptides

We next assessed whether the c-Src SH3 *H*3_2_ crystal lattice is compatible with typical PxxP peptide binding, to ask if this crystal form might be amenable to in-crystal binding studies [53]. SH3 domains bind typical PxxP-containing peptides via a binding pocket between the RT- and n-Src loops. This binding site can be broken up into three sub-pockets: two shallow Proline-binding (xP) pockets that accommodate the minimum PxxP binding motif and a third specificity site that binds residues in an extended peptide (Figure 5A) [10]. We superposed a co-crystal structure of the Src SH3 domain bound to the VSL12 peptide (PDB ID: 1rtz, unpublished, R.M.S.D. 1.1 - 1.4 Å over 56 equivalent Cα positions) onto each chain of our crystal structure and examined the positions of the modelled PxxP peptide position. Within the asymmetric unit, PxxP peptide binding to each of the four chains is compatible with the SH3 domain arrangement (Figure 5B). Next, we extended the asymmetric unit by generating symmetry mates using crystal symmetry and examined the symmetry-related lattice contacts. We observe two distinct lattice contact arrangements between chains in the asymmetric unit and symmetry-related molecules. First, chain A interacts with symmetry-related chain D (Figure 5C), and this lattice contact arrangement is shared between chain B and symmetry-related chain C (Figure S4A). When modeling a PxxP peptide bound to the SH3 domain, we observe that his lattice contact site sterically clashes with the predicted position of the PxxP peptide. Second, chain C interacts with symmetry-related chain C (Figure 5D) which is shared between chain D and symmetry-related chain D (Figure S4B), but distinct from that observed for chains A and B (Figure 5C). In this arrangement, PxxP peptide binding to the typical binding site is also prevented by lattice contacts. We thus conclude that that standard PxxP peptide binding to the SH3 domain at the xP and specificity sites is incompatible with formation of the *H*3_2_ crystal lattice since these sites in the SH3 domain are involved in crystal lattice contacts.

**Figure 5.**
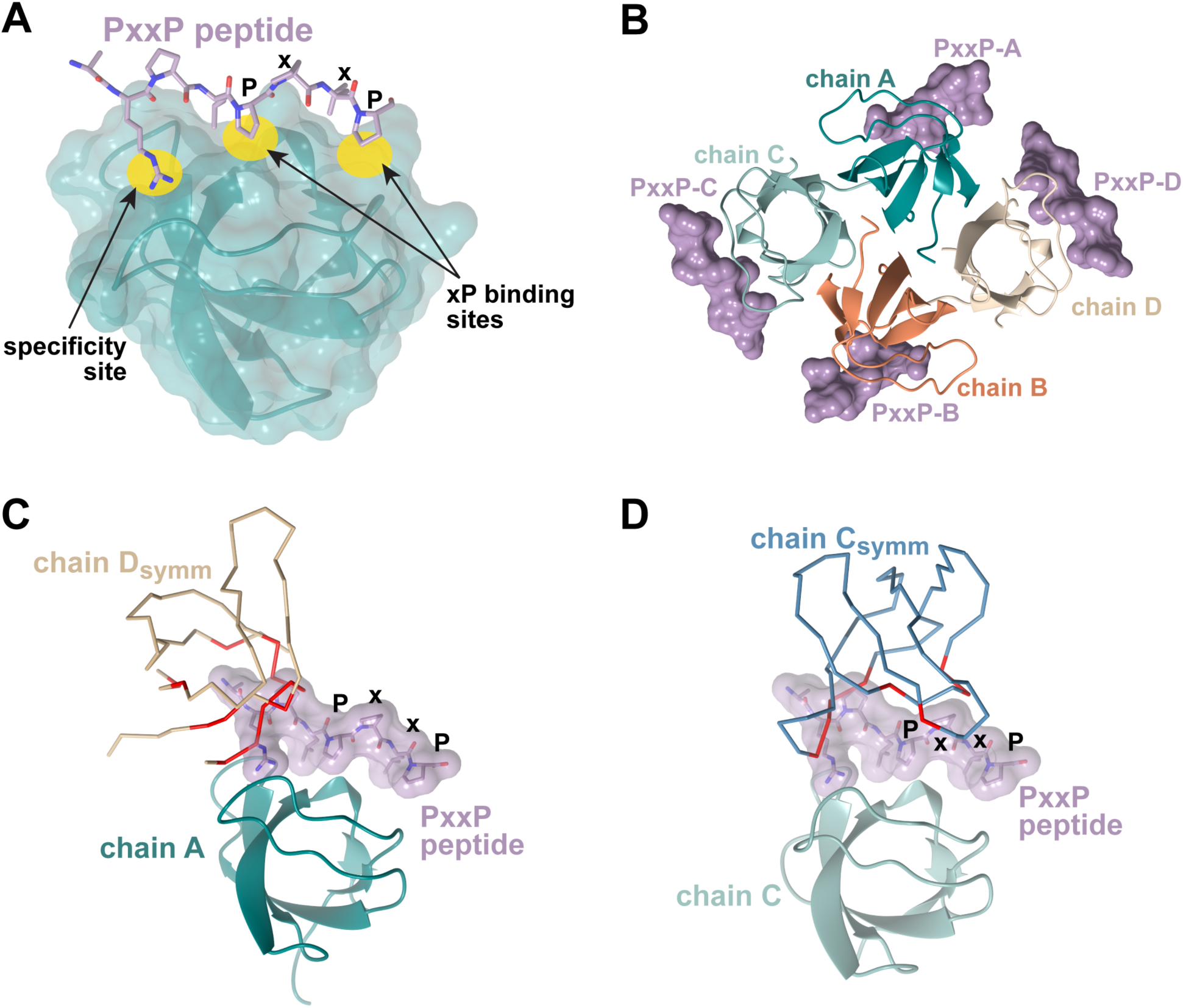
**Modeling PxxP binding onto c-Src SH3 domains in *H*3_2_ space group.** (A) Co-crystal structure of Src SH3 domain (teal, drawn in ribbon and semi-transparent surface) bound to the VSL12 peptide (in purple) (PDB ID: 4rtz, unpublished). Peptide binding sites on SH3 are highlighted in yellow and labeled. (B) Asymmetric unit of c-Src SH3 *H*3_2_ crystal structure. The PxxP peptide (purple surface) is modeled onto each chain by superposition of the complex from (A) with each individual chain of the asymmetric unit, with R.M.S.D. values ranging from 0.7 - 1.0 over 57 equivalent Cα positions. (C) and (D) Modeling reveals that the predicted PxxP binding sites are blocked by symmetry-related molecules in the *H*3_2_ crystal form. Residues in the symmetry mates (Chain D_symm_ (C) and Chain C_symm_ (D)) with predicted steric overlaps with the modeled PxxP peptide are colored red on the Cα trace (calculated in MolProbity [67]). (C) c-Src SH3 chain A (teal ribbon) and its symmetry-related chain D (D_symm_, tan Cα trace). Chain B shares a similar interface with a symmetry-related chain C (Figure S4A). (D) c-Src SH3 chain C (light cyan ribbon) and symmetry-related chain C (C_symm_, grey-blue). Chain D shares a similar interface with a symmetry-related chain D (Figure S4B).

## 4. DISCUSSION AND CONCLUSION

In this study we describe a new crystal form for the c-Src SH3 domain protein that is dependent on interchain metal-mediated coordination and unique crystal contacts and expands the crystallization chemical space and lattice types for this important regulatory domain. This is achieved due to the native N-terminus of the isolated c-Src SH3 protein creating a previously unobserved binding site for Ni^2+^ ions. This finding offers potential benefits in biotechnology. For example, proteins in the environment of a crystal lattice are used as targets in drug discovery like crystal fragment-based screens [54]. The ability to obtain diverse lattice types for different protein targets affords a wider scope of available binding sites for fragment based screening [55].

In this example, metal binding facilitates protein crystallization. Multiple systematic approaches to improve the propensity of purified proteins to form well-ordered crystals for X-ray diffraction studies, also called “crystal lattice engineering,” have been proposed. For example, protein modification tools including mutational surface entropy reduction [56] or enzymatic lysine methylation [57] have been described as approaches to promote protein crystal lattice formation. Other approaches, sometimes termed synthetic (or mutational) symmetrization, involve engineered surface amino acid mutations to promote protein self-multimerization in a symmetric arrangement. This includes mutations to drive disulfide bond formation [58, 59], surface leucine zippers [60], and metal coordination [61, 62].

Here, we provide evidence of metal-mediated symmetrization in the absence of the additional step of creating synthetic (or mutational) sites. For the c-Src SH3 domain, this metal-mediated crystallization instead uses native amino acid residues: an N-terminal glycine residue that is the result of engineering of the protein expression construct, and a surface-exposed histidine in a second molecule. This raises the possibility of metal-mediated symmetrization in otherwise recalcitrant protein samples, as has been observed previously (for example, [63, 64]). Thus, inclusion of metal ions during initial crystallization screening may help proteins self-associate into a crystal lattice. Common divalent metal salts include NiCl_2_, CdCl_2_, CoCl_2_, and CuSO_4_, which are readily accessible in many protein crystallography labs. Additionally, the absorption edge wavelengths of these common metal ions are readily achievable at standard synchrotron beamlines, which may allow for anomalous signal to be recorded and used for metal ion placement in the three-dimensional model as well as phasing for macromolecular structure solution [37, 65].

Lastly, our study demonstrates that continued studies of the SH3 domain can yield new results that expand the current knowledge of this important modular protein domain. Currently, a search of the term “SH3 domain” in the PDB yields over 6,000 entries for X-ray crystal structures, highlighting the abundance of structural information for this ubiquitous domain. SH3 domain structures were once referred to as the “darlings of the structural biology set” [24] due to their prominence. Importantly, the function and activity of SH3 domains was first elucidated, in part, by structural biology studies which yielded three dimensional models of these domains. For example, three dimensional structures helped uncover the site of PxxP ligand binding, of specificity determinants, of autoinhibition within the Src family kinases, and atypical binding with non-PxxP binding partners to alternative sites on the SH3 domain [9, 17]. Such sustained structural studies of SH3 domains have proven beneficial. The three dimensional structure of a Src family kinase SH3 domain was first observed in 1992-1993 [6, 7], and Src SH3 continues to be a current target of structural studies (for example [66]).

## Supporting information

Plasmid sequence

Plasmid map

## ACKNOWLEDGEMENTS

Anton Bennett and Sravan Perla are thanked for helpful discussions. This research used Center for BioMolecular Structure (CBMS) beamline 17ID-2 (FMX) of the National Synchrotron Light Source II, a U.S. Department of Energy (DOE) Office of Science User Facility operated for the DOE Office of Science by Brookhaven National Laboratory under Contract No. DE-SC0012704. CBMS is primarily supported by the National Institutes of Health, National Institute of General Medical Sciences (NIGMS) through a Center Core P30 Grant (P30GM133893), and by the DOE Office of Biological and Environmental Research (KP1607011). Data were collected using the beamtime obtained through NE-CAT BAG proposal #311401. We thank Alexei Soares for assistance with data collection. X.C. is a 2024 recipient of a Yale College Dean’s Research Fellowship. O.S.F. is supported by NIH R35GM150644. This work is supported by NIH R01GM102262 to T.J.B.

## DATA AVAILABILITY

X-ray crystal structure model coordinates have been deposited in the RSCB Protein Data Bank (PDB) under accession code 9OFX (https://doi.org/10.2210/pdb9OFX/pdb). X-ray diffraction images are available online at SBGrid Data Bank under dataset number 1158 (10.15785/SBGRID/1158). The expression plasmid map and full plasmid sequence are provided as supplementary materials.

