## Supplementary information for "Nickel binding to c-Src SH3 domain facilitates crystallization"

**Research Article**

**Table S1. Macromolecule production information.**

| | |
|---|---|
| Source organism | *Homo sapiens* |
| DNA source | cDNA |
| Expression vector | pET-28a(+) |
| Plasmid construction method | PCR and restriction-ligation |
| Forward primer (NheI and TEV site) | 5' - GGATTCCATATGGAGAACCTGTACTTTCAA GGCGTGACCACCTTTGTGGCCC - 3' |
| Reverse primer (STOP codon and XhoI) | 5' - CCGCTCGAGCTAGGAGGGCGCCACATAGTTG - 3' |
| Expression host | *E. coli* Rosetta(DE3) |
| Expression details | Induced with 0.2 mM IPTG at $OD_{600}$ = 0.8, 18°C, 20 hr |
| Complete amino acid sequence of the protein produced | MGSSHHHHHHSSGLVPRGSHMENLYFQGVTTFVALYDY ESRTETDLSFKKGERLQIVNNTEGDWWLAHSLSTGQT GYIPSNYVAPS |
| Complete amino acid sequence of the protein after TEV proteolysis | 85 - GVTTF VALYDYESRT ETDLSFKKGE RLQIVNNTEG DWWLAHSLST GQTGYIPSNY VAPS - 143 |

**Table S2. Crystallization information.**

| | |
|---|---|
| Method | Hanging drop vapor diffusion |
| Plate type | VDX™ Plate with sealant (Hampton Research) |
| Temperature (°C) | 25º C (room temperature) |
| Protein concentration | 1.8 mM, 11.9 mg/mL |
| Buffer composition of protein solution | 150 mM NaCl, 20 mM Tris pH 7.5 |
| Composition of reservoir solution | 1.7 M Ammonium Sulfate, 5 mM $NiCl_2$, 10% glycerol, 0.1 M HEPES pH 7.5 |
| Volume and ratio of drop | 2 µL drop, 1:1 v/v protein to reservoir ratio |
| Volume of reservoir | 500 µl in VDX plate (Hampton Research) |
| Composition of the cryopreservative | 1.7 M Ammonium Sulfate, 5 mM $NiCl_2$, 20% glycerol, 0.1 M HEPES pH 7.5 |
| Drop setting | Manual |
| Seeding | No |

**Table S3. Additional Data collection and processing statistics.**

| | | | | |
|---|---|---|---|---|
| Diffraction source | BNL NSLS-II 17-ID-2 FMX | | | |
| Detector | EIGER 16M | | | |
| Numver of crystals merged | 4 | | | |
| Individual crystals | 1 | 2 | 3 | 4 |
| Temperature (K) | 100 | 100 | 100 | 100 |
| Crystal to detector distance (mm) | 200 | 200 | 200 | 200 |
| Total rotation range (°) | 240 | 240 | 240 | 240 |
| Rotation per image (°) | 0.2 | 0.2 | 0.2 | 0.2 |
| Exposure time per image (s) | 0.01 | 0.01 | 0.01 | 0.01 |
| Space group | $H3_2$ | $H3_2$ | $H3_2$ | $H3_2$ |
| Unit cell dimensions | | | | |
| a (Å) | 63.760 | 63.976 | 63.976 | 63.867 |
| b (Å) | 63.760 | 63.976 | 63.976 | 63.867 |
| c (Å) | 271.669 | 272.214 | 272.816 | 272.628 |
| α, β, γ (°) | 90, 90, 120 | 90, 90, 120 | 90, 90, 120 | 90, 90, 120 |
| Mosaicity (°) | 0.09 | 0.06 | 0.07 | 0.06 |
| Anomalous resolution limit (Å)[+] | 2.63 | 2.40 | 2.16 | 2.21 |

[+] Anomalous signal detected in Aimless (Evans and Murshudov, 2013) where $CC_{anom} < 0.15$

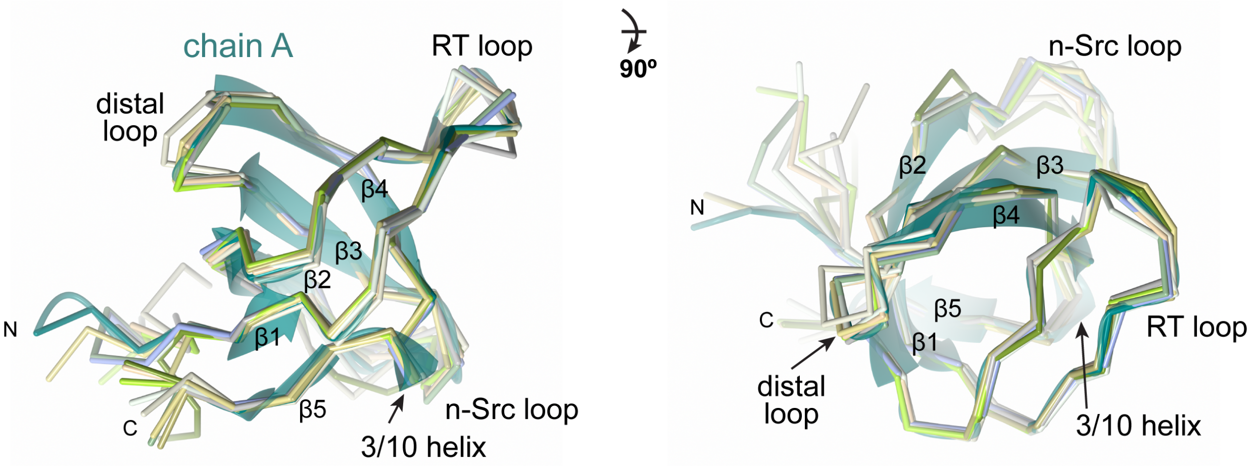


**Figure S1. Superposition of 10 representative c-Src SH3 domain structures.**
Chain A from the crystal structure in this study shown in dark teal ribbons. Superposed are a representative set of c-Src SH3 domain crystal structures from the PDB. Secondary structure elements and loops are labelled. The view in the right panel is related to the left view by a 90° rotation toward the reader about the x-axis. The R.M.S.D. of superposition ranges from 0.55 - 1.09 Å over 58 equivalent Cα positions for the ten representative structures shown. Superpositions were performed using the DALI server (Holm and Rosenstrom, 2010). The ten representative structures are as follows: PDB: 4jz4 (Bacarizo et al., 2014), 4omo (Bacarizo et al., 2014), 4rtx (unpublished), 4rtz (unpublished), 6c4s (Kall et al., 2019), 4hxj (Xiao et al., 2013), 4hvw (Bacarizo and Camara-Artigas, 2013), 5eca (unpublished), 7a3c (unpublished), 7a32 (unpublished).

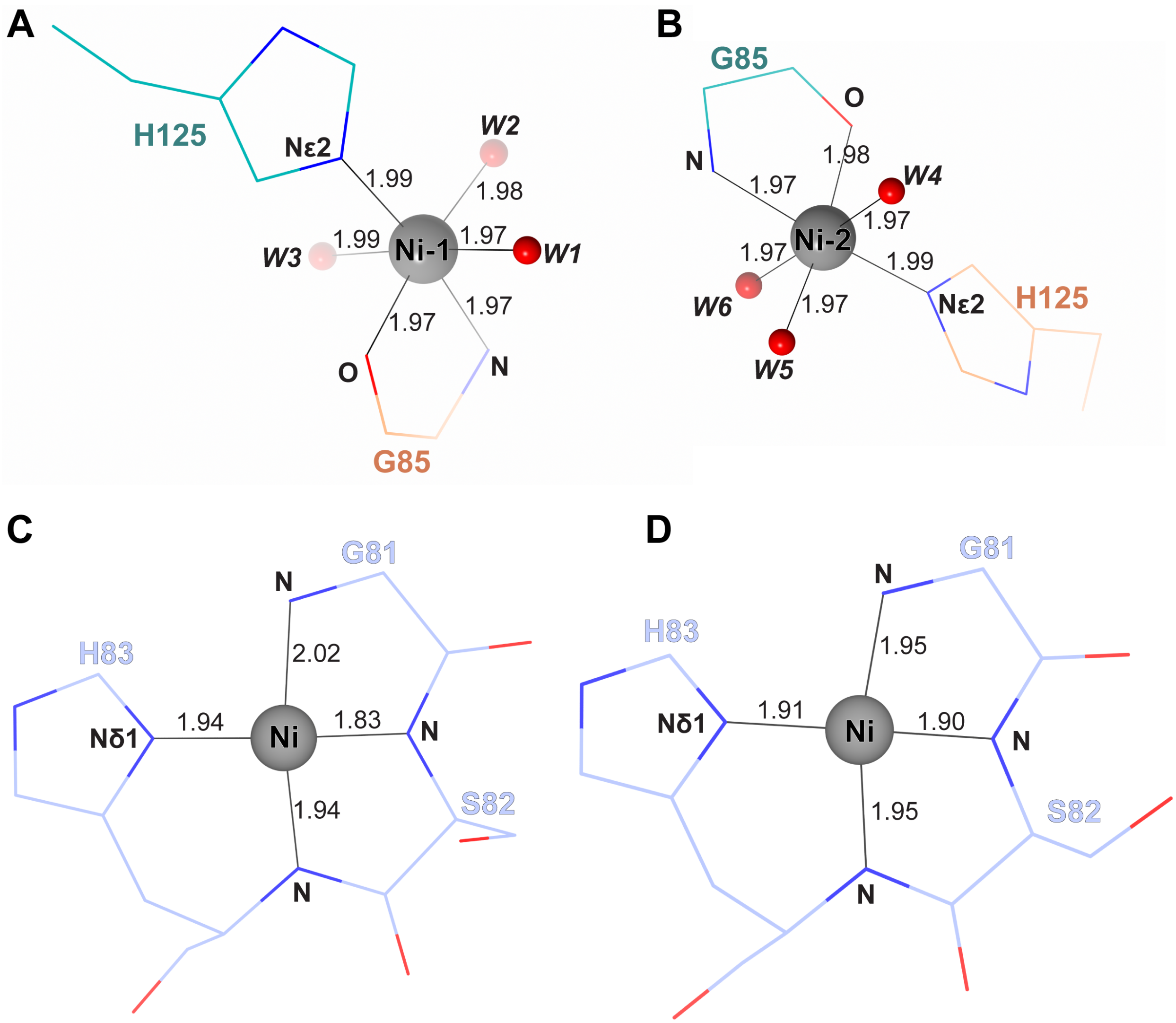


**Figure S2. Metal-ligand bond distances.**
(A) c-Src SH3 structure in $H3_2$ crystal form with octahedral coordination of nickel (Ni-1) by His-125 of chain A (teal), Gly-85 of chain B (orange), and three water molecules (red spheres), showing the metal-ligand distance for each bond.
(B) Similar to part (A), the octahedral coordination of nickel (Ni-2) by Gly-85 of chain A (teal), His-125 of chain B (orange), and three water molecules (red spheres).
In both panels, the water molecules are numbered according to the final PDB model. These nickel-ligand distances are also listed in Table 5.
(C) and (D) Metal-ligand bond distances in classic ATCUN motif bound to nickel in square planar geometry in the Src SH3 structure (PDB: 4jz4 (Bacarizo et al., 2014)) chains A (panel C) and B (panel D). All bond distances are reported by CheckMyMetal server (Gucwa et al., 2023).

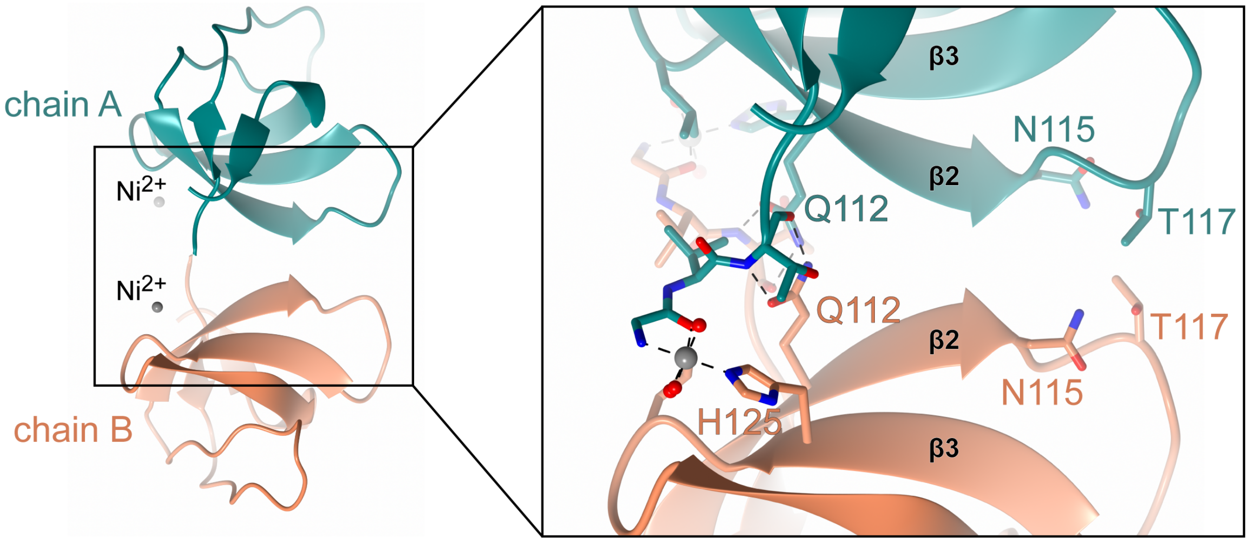


**Figure S3. Additional details of the interface between chains A and B in the asymmetric unit.**
Left panel: Chains A (dark teal) and B (orange) as ribbons and two $Ni^{2+}$ ions (grey spheres) in the asymmetric unit. The view is related to Figure 2c by a 90° rotation to the left on the y-axis.
Right panel: Zoom-in view shows the details of the interface including the sidechains of Asn-115 and Thr-117 in both chains which participate in van der Waals interactions between the chains.

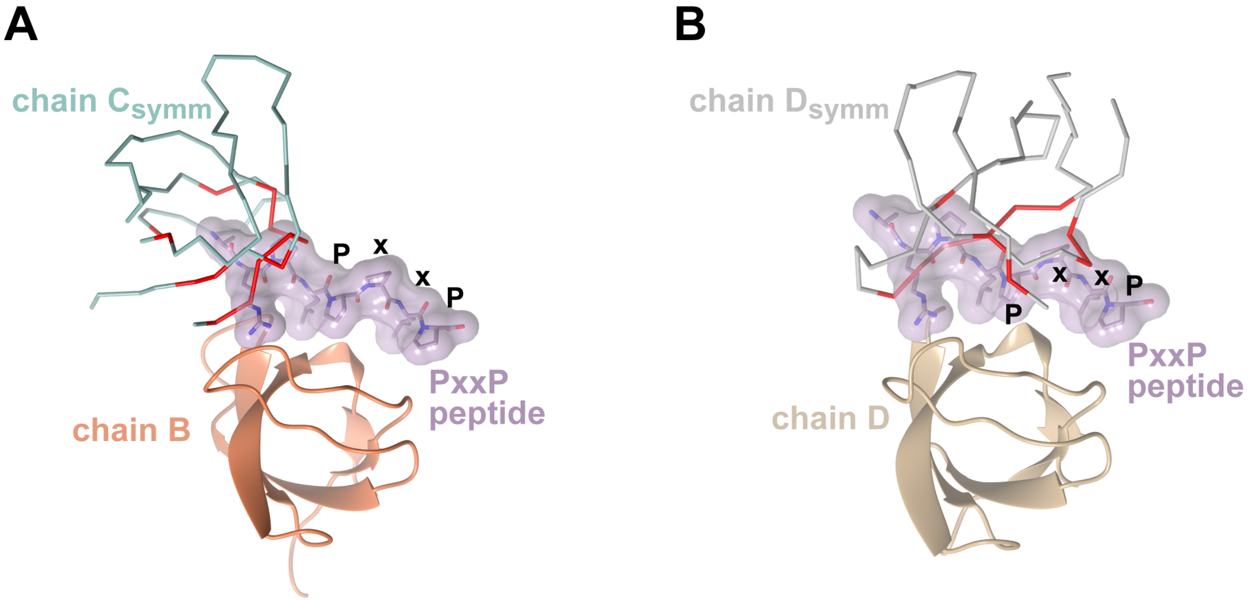


**Figure S4. Predicted PxxP peptide binding is blocked by symmetry-related molecules.**
The predicted PxxP binding sites are blocked by symmetry-related molecules in the $H3_2$ crystal form. Related to Figure 5.
(A) c-Src SH3 chain B (orange ribbon) and its neighboring lattice partner chain $D_{symm}$ (light cyan Cα trace). This interface is similar to the one shown in Figure 5C.
(B) Chain D (tan ribbon) and its neighboring lattice partner chain $D_{symm}$ (grey Cα trace). This interface is similar to the one shown in Figure 5D. In (A) and (B), residues in the symmetry mates (chain $D_{symm}$ (A) and Chain $D_{symm}$ (B)) with predicted serious steric overlaps with the modeled PxxP peptide are colored red on the Cα traces (calculated in MolProbity (Chen et al., 2010)).
