## Supplementary figures and images for "Nickel binding to c-Src SH3 domain facilitates crystallization"

### Plasmid map

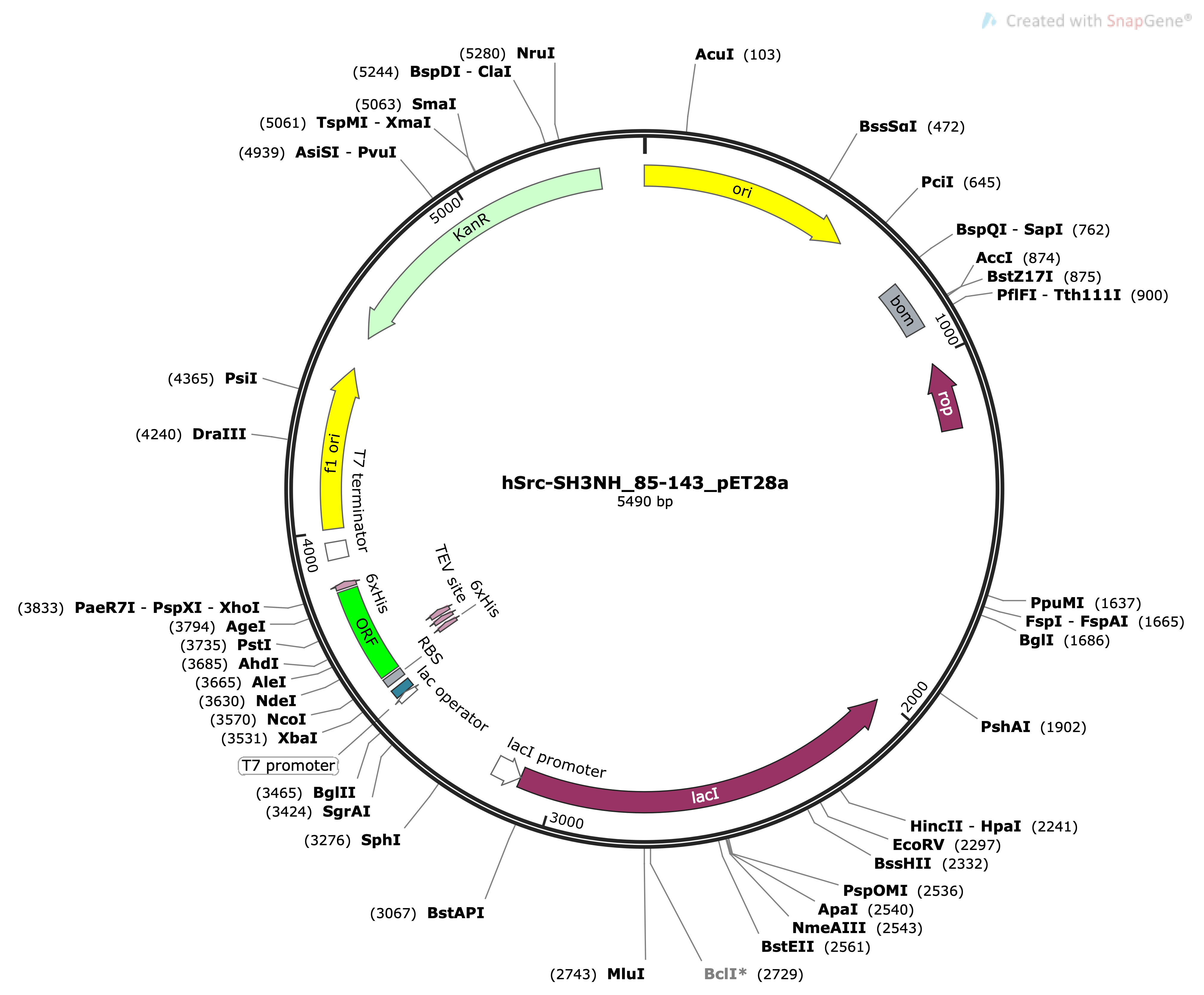
